# An examination of multivariable Mendelian randomization in the single sample and two-sample summary data settings

**DOI:** 10.1101/306209

**Authors:** Eleanor Sanderson, George Davey Smith, Frank Windmeijer, Jack Bowden

## Abstract

**Background:** Mendelian Randomisation (MR) is a powerful tool in epidemiology which can be used to estimate the causal effect of an exposure on an outcome in the presence of unobserved confounding, by utilising genetic variants that are instrumental variables (IVs) for the exposure. This has been extended to Multivariable MR (MVMR) to estimate the effect of two or more exposures on an outcome.

**Methods/Results:** We use simulations and theory to clarify the interpretation of estimated effects in a MVMR analysis under a range of underlying scenarios, where a secondary exposure acts variously as a confounder, a mediator, a pleiotropic pathway and a collider. We then describe how instrument strength and validity can be assessed for an MVMR analysis in the single sample setting, and develop tests to assess these assumptions in the popular two-sample summary data setting. We illustrate our methods using data from UK biobank to estimate the effect of education and cognitive ability on body mass index.

**Conclusion:** MVMR analysis consistently estimates the effect of an exposure, or exposures, of interest and provides a powerful tool for determining causal effects in a wide range of scenarios with either individual or summary level data.

## Key Messages

- Multivariable Mendelian randomisation (MVMR) has been introduced as a technique for estimating the causal effect of multiple exposure variables on a health outcome with two-sample summary data. We build on this work by clarifying how MVMR should be applied with individual level data and two-sample summary data, in order to conform with established econometric theory for multivariable two-stage least squares analysis.
- Instrument strength and validity should be assessed in the single sample MVMR setting using the Sanderson-Windmeijer F statistic and the Sargan test.
- We develop a generalised version of Cochran’s Q statistic to test for instrument strength and validity in the two-sample summary data setting. However, these tests require knowledge of the covariance between the effects of the genetic variants on each exposure.
- If the covariance between the effect of the genetic variants on each exposure can be either: (i) estimated from individual data; (ii) assumed to be zero, or; (ii) fixed at zero by using non-overlapping samples for each exposure GWAS, then our proposed summary data Q statistics will give a good approximation of the true (individual level data) result.
- The causal effect estimated by Mendelian Randomisation and Multivariable Mendelian Randomisation can differ.MR estimates the total causal effect of the exposure on the outcome, whereas MVMR estimates the *direct causal* effect of each exposure on the outcome.

## Introduction

In many scenarios where we wish to estimate the causal effect of an exposure X on an outcome Y, a conventional regression analysis can be misleading, as the observational association between the two variables could easily be affected by unobserved confounding. If genetic variants – usually single nucleotide polymorphisms (SNPs) - are available which reliably predict the exposure variable but do not have an effect on the outcome through any other pathway, then they are valid instrumental variables (IVs) and can be used in a Mendelian randomization (MR) analysis to obtain unconfounded estimates of the effect of the exposure on the outcome, as illustrated in Figure 1.

**Figure 1:**
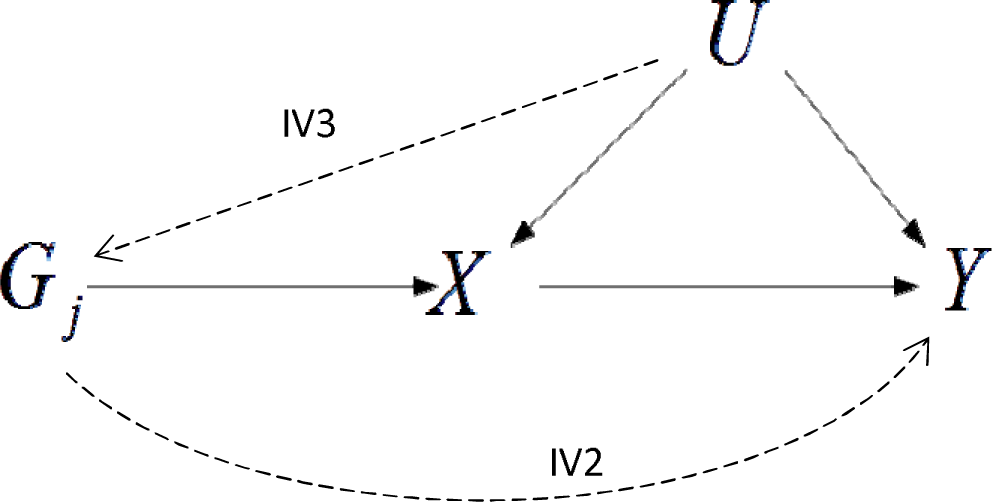
Hypothesised relationship between genetic variant G_j_, modifiable exposure X and outcome Y in the presence of an unobserved confounder, denoted by U. The line from G_j_ to X represents instrumental variable assumption IV1. Dashed lines represent potential violations of the instrumental variable assumptions IV2 and IV3.

In many scenarios we may wish to estimate the effect of multiple exposures on the outcome using MR analysis, for example; because we believe these exposures to be closely related or because we believe one exposure may mediate the relationship between the exposure of primary interest and the outcome. This can be done with Multivariable MR (MVMR) where a set of genetic variants is used to predict a set of exposure variables. However, careful consideration needs to be given in such an analysis to exactly what relationship is being estimated and how the IV assumptions required for MR analysis apply to a MVMR analysis. In this paper we build on previous work developing MVMR methods with two-sample summary level data (1, 2) and fully explain how MVMR can be implemented with either individual level or two-sample summary level data, exactly what is being estimated in a MVMR analysis and how the IV assumptions required for MR analysis translate to MVMR analysis. We describe existing tests that can be used to test the IV assumptions with individual level data and add to the previous literature on MVMR by developing new methods to identify potential violations of the IV assumptions with two-sample summary level data.

### Mendelian Randomisation

To state the IV assumptions more formally with reference to Figure 1: For a single SNP *G_j_* to be a valid IV it must be:

IV1: associated with the exposure *X* (the “relevance” assumption);

IV2: independent of the outcome *Y* given the exposure *X*, (the “exclusion restriction”); and

IV3: independent of all (observed or unobserved) confounders of *X* and *Y*, as represented by U (the “exchangeability” assumption);

If IV1-IV3 are satisfied for a set of SNPs *G* = (*G*_l_,…, *G*_L_), then traditional IV methods can be employed to reliably test for a causal effect of *X* on *Y* using *G, X* and *Y* alone, without any attempt to adjust for *U* at all. For example, suppose the variables G, X, U and Y are linked via the following models:

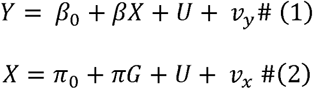

Here *v_x_* and *v_y_* represent independent error terms, *π* represents the parameter vector *π*_l_,…, *π_L_*, and *β* is the true causal effect of *X* on *Y* we wish to estimate. We will assume throughout this paper that (*G*_l_,…, *G_L_*) are mutually uncorrelated (by design). A naïve regression of *Y* on *X* will not yield a consistent estimate for *β* because the explanatory variable in the regression, *X*, is correlated with *U*. However, two-stage least squares (TSLS) estimation, performed by regressing *Y* instead on *X̂* the predicted value of *X* from a regression of *X* on *G – will* yield a consistent estimate for *β*, because *X̂* is independent of u.(3, 4)

TSLS relies on individual level data, but the sharing of such data is often impractical. In recent years it has become much more common to attempt MR analyses using summary data estimates of SNP-exposure and SNP-outcome associations gleaned from two independent but homogeneous study populations. The SNPs in question are usually identified as genome-wide significant ‘hits’ in distinct genomic regions via a genome wide association study (GWAS) for the exposure. This is referred to as ‘two-sample summary data MR’.

Let *π_j_* and Γ_*j*_ represent the true association for SNP *G_j_* in *G* with the exposure and the outcome respectively. From models (1) and (2) we can link the *j*’th SNP outcome association to the *j*’th SNP exposure association via the model

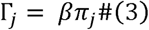

It follows that the Wald estimator 
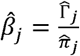
, is a consistent estimate for *β*. (5, 6) Where Γ̂*_j_* and *π̂_j_* are estimates from OLS estimation of;

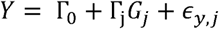

And

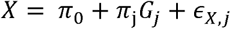

When the SNPs are uncorrelated, taking an inverse variance weighted (IVW) average of the ratio estimates will yield an overall estimate for *β*, *β̂_IVW_*, that closely approximates the TSLS estimate that would have been obtained if individual level data were available.(7)

### Detecting ‘weak’ instruments and ‘invalid’ instruments in MR

If assumptions IV1 – IV3 are fulfilled for all SNPs in *G*, and linear models (1)-(2) hold, then either a TSLS or IVW analysis (with uncorrelated SNPs) will consistently estimate the causal effect.(8–10) In order to satisfy IV1, the SNPs in *G* should strongly predict the exposure *X*. This can be quantified using the F-statistic from the first stage regression of *X* on *G*. Using instruments that are jointly only weakly associated with the exposure (i.e. which have a small F-statistic) will result in weak instrument bias.(11)

Secondly, SNPs should not exert a direct effect on *Y*, i.e. they should not affect *Y* other than through *X*. Any such effect would represent a violation of IV2. Horizontal pleiotropy, where the genetic variants used as instruments have an effect on the outcome that is not through the exposure of interest is a violation of the exclusion restriction and could easily be responsible for such a violation in the MR setting.(10, 12–14) The SNPs should also not be confounded by any variables that also influence the outcome. Any confounding of this nature would be a violation of assumption IV3. A violation of either assumptions IV2 or IV3 is likely to lead to bias and potentially erroneous conclusions in both the TSLS and IVW estimates.(4) The presence of potential pleiotropic effects can be evaluated using the Sargan test(15, 16) using individual level data and Cochran’s Q statistic(17–19) using summary data. The causes and consequences of pleiotropy in MR are described in detail elsewhere.(1, 9, 10, 13, 14, 20)

In addition to assumptions IV1 – IV3 there are additional assumptions and considerations that apply to all instrumental variable estimation, including MR and MVMR. These included the assumptions of linearity and homogeneity which are in many settings required for obtaining a point estimate of the causal effect. However, if this assumption is violated the causal null will still be respected and it will still be possible to identify whether the exposures are causally associated with the outcome.(21–23) Throughout this analysis we assume linearity of the relationship between the exposures and the outcome, however if this assumption did not hold the same issues would apply to MVMR as apply in MR analysis which are discussed in detail elsewhere.(24, 25)

## Multivariable Mendelian randomization

MR can be extended to estimate the effect of multiple exposure variables on an outcome(1) and is particularly useful in cases where a standard MR analysis would fail due to violation of assumptions IV2-3. It is also useful in cases where two or more correlated exposures are of interest and may help to understand if both exposures exert a causal effect on the outcome, or if one in fact mediates the effect of the other on the outcome(26, 27). ‘Multivariable MR’ (MVMR) requires a set of SNPs, *G*, which are associated with the exposure variables but do not affect the outcome other than through these variables. In the same way as standard (single variable) MR, these SNPs can be used to predict each of the exposure variables in the model and these predicted values can be used to estimate the effect of the exposures on the outcome in a multivariable regression analysis. The setup for MVMR is illustrated for an analysis involving two exposure variables (*X*_l_ and *X*_2_) in Figure 2. The arrows linking *X*_l_ with *X*_2_, and *X*_2_ with *Y* have been left bi-directional to acknowledge the fact that many underlying causal relationships are possible. That is, they could point in either direction or be completely absent. Indeed, many of these options will be subsequently explored.

**Figure 2:**
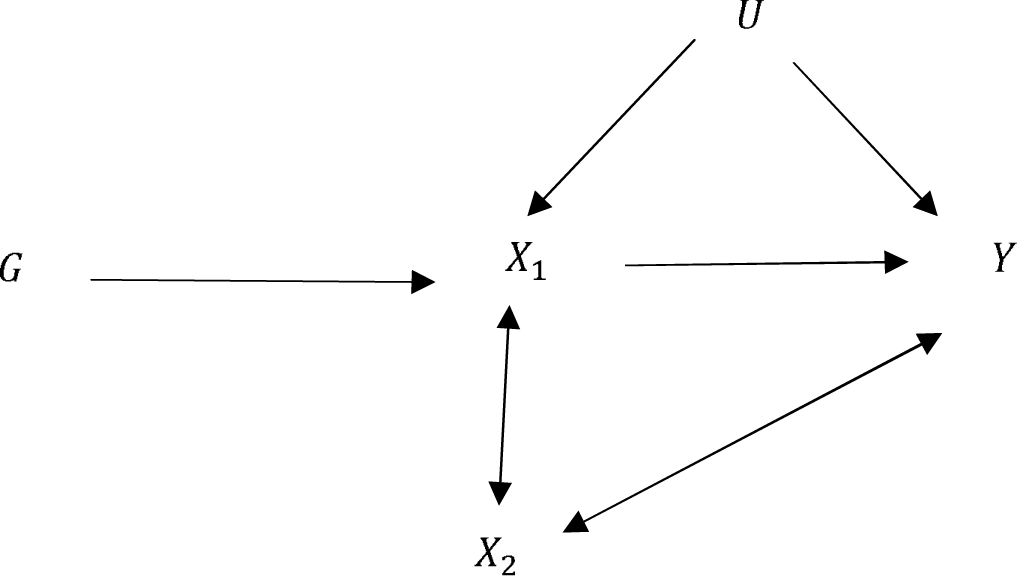
Hypothesised relationship between genetic variant(s) G, modifiable exposures, X_l_, X_2_ and outcome Y in the presence of unobserved confounder U. Bi-directional arrows represent possible violations of the IV assumptions induced by X_2_ that are explored in this paper.

Although it is the simplest possible MVMR setting, two exposures suffice to illustrate all the scenarios and ideas described in this paper. From Figure 2, we can write the following general model linking *Y, X*_l_, *X*_2_ and *U*:

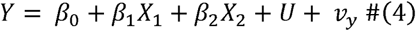

For example, suppose that *X*_1_ and *X*_2_ are in fact independent given G (so there is **no** arrow in Figure 2 between *X*_1_ and *X*_2_) and *X*_2_ affects *Y* independently of *X*_1_ (so that there is a **direct arrow** from *X*_2_ to Y). If true, then models (5) and (6) for *X*_1_ and *X*_2_ would, jointly with (4), describe the data:

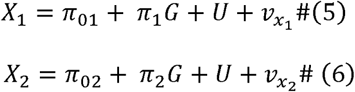

The purpose of an MVMR analysis is to determine the direct causal effect of both *X*_1_ and *X*_2_ on the outcome *Y*, when conditioned on one another. Without loss of generality we will focus primarily on the effect of *X*_1_ (and the parameter *β*_1_ with the direct effect of *X*_2_ on *Y* denoted by *β*_2_ being of secondary importance.

With individual level data TSLS can be implemented with multiple exposure variables, regressing each exposure on the full set of SNPs to yield genetically predicted estimates for *X*_1_ and *X*_2_. The outcome Y can then be regressed on these predicted estimates for *X*_1_ and *X*_2_ jointly to obtain consistent estimates of *β*_l_ and *β*_2_. This can be conducted by simply using the *ivreg2* command in Stata or *ivpack* in R.

In the two sample summary data setting, Burgess and colleagues(1, 2) show how MVMR can be implemented using summary data estimates of the association between SNP *J*(out of *L*) and: the outcome, Γ̂_*j*_; exposure *X*_1_, *π̂*_l*j*_; and exposure *X*_2_, *π̂*_2*j*_, by fitting the following model:

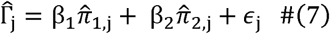

This is a straightforward generalization of the IVW estimation framework.

### Important considerations

To conduct an MVMR analysis it is necessary to have at least as many genetic instruments as there are exposures to be instrumented in the model, this is true regardless of whether single sample or two sample summary data are used. It is possible to include genetic instruments that are associated with more than one exposure variable, providing all of those exposure variables are included in the estimation. Instruments must not, however, exert a direct effect on the outcome, except through the included exposures. There is no benefit to excluding instruments that are only strongly associated with one exposure, as this will lead to a loss of precision in the estimates obtained. This also avoids any potential bias that could arise due to selecting instruments based on their strength.(11)

## What quantities do MR and MVMR estimate?

MR and MVMR target different causal effects of the exposure on the outcome. In general, MR estimates the total effect of the exposure on the outcome, whereas MVMR estimates the direct effect of each exposure on the outcome.

For example, if Figure 3 describes the truth, the total effect of exposure *X*_1_ on the outcome is the effect of *X*_1_ on the outcome *Y* directly plus the effect of *X*_1_ on *Y* via *X*_2_, and is equal to *β*_l_ + *αβ*_2_. The direct effect of the exposure *X*_1_ on the outcome *Y* is the effect *X*_1_ has on *Y* not via any other exposure variables included, and so is equal to *β*_l_. Whether or not these effects differ in general depends on the underlying relationship between the exposures and between each exposure and the outcome. If there is no effect of *X*_1_ on *X*_2_ or of *X*_2_ on *Y*, i.e. either *α* or *β*_2_ is equal to zero, these effects will be the same.

**Figure 3:**
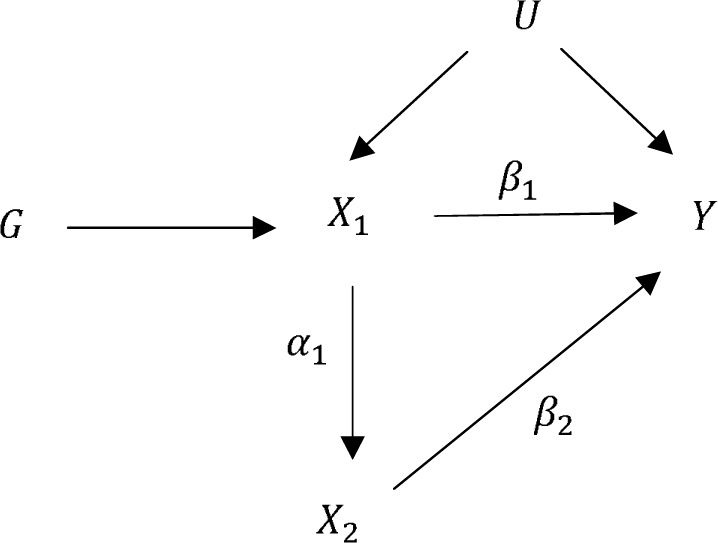
Illustration of the direct effect and total effect of X_1_ on the outcome Y.

To highlight the potential differences between MR and MVMR, and the potential benefits of MVMR, we now consider the application of MVMR to four different scenarios which are commonly encountered, or at least suspected, in epidemiological studies

. Each of these scenarios represents a situation where conventional univariable MR would produce consistent results, given the correct set of SNPs, but where MVMR may estimate a different causal effect and provide benefits when in fact some of the SNPs may have effects on more than one exposure (and thus making them invalid instruments for a univariable MR analysis). In the first scenario *X*_2_ is a **confounder** of the relationship between *X*_1_ and *Y*. That is, there is a direct causal path from *X*_2_ to *X*_1_ and from *X*_2_ to *Y*. Along with model (4), model (6) above and (8) below underlie the individual level data:

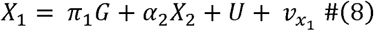

In the second scenario *X*_2_ is a **collider** of the relationship between *X*_1_ and *Y*. That is, there is a direct causal path from *X*_1_ to *X*_2_ and from *Y* to *X*_2_. When an exposure and outcome both influence another variable, controlling for that variable in conventional analysis will introduce bias into the observed association between the exposure and the outcome.(28) This form of bias can also be understood as a form of selection bias which would occur if inclusion in the sample was dependent on the value of *X*_2_.(29) Along with model (4) (with *β*_2_ set to 0), model (5) above and (9) below are used to generate the individual level data:

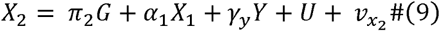

In the third scenario is an independent **pleiotropic** pathway from to ․ This corresponds to the scenario first described in the previous section. Along with model (4), models (5) and (6) above are used to generate the individual level data.

In the fourth scenario is a **mediator** of the relationship between and ․ Along with model (4), model (5) above and (10) below are used to generate the individual level data:

Each of these scenarios are shown in Figure. 4.

**Figure 4:**
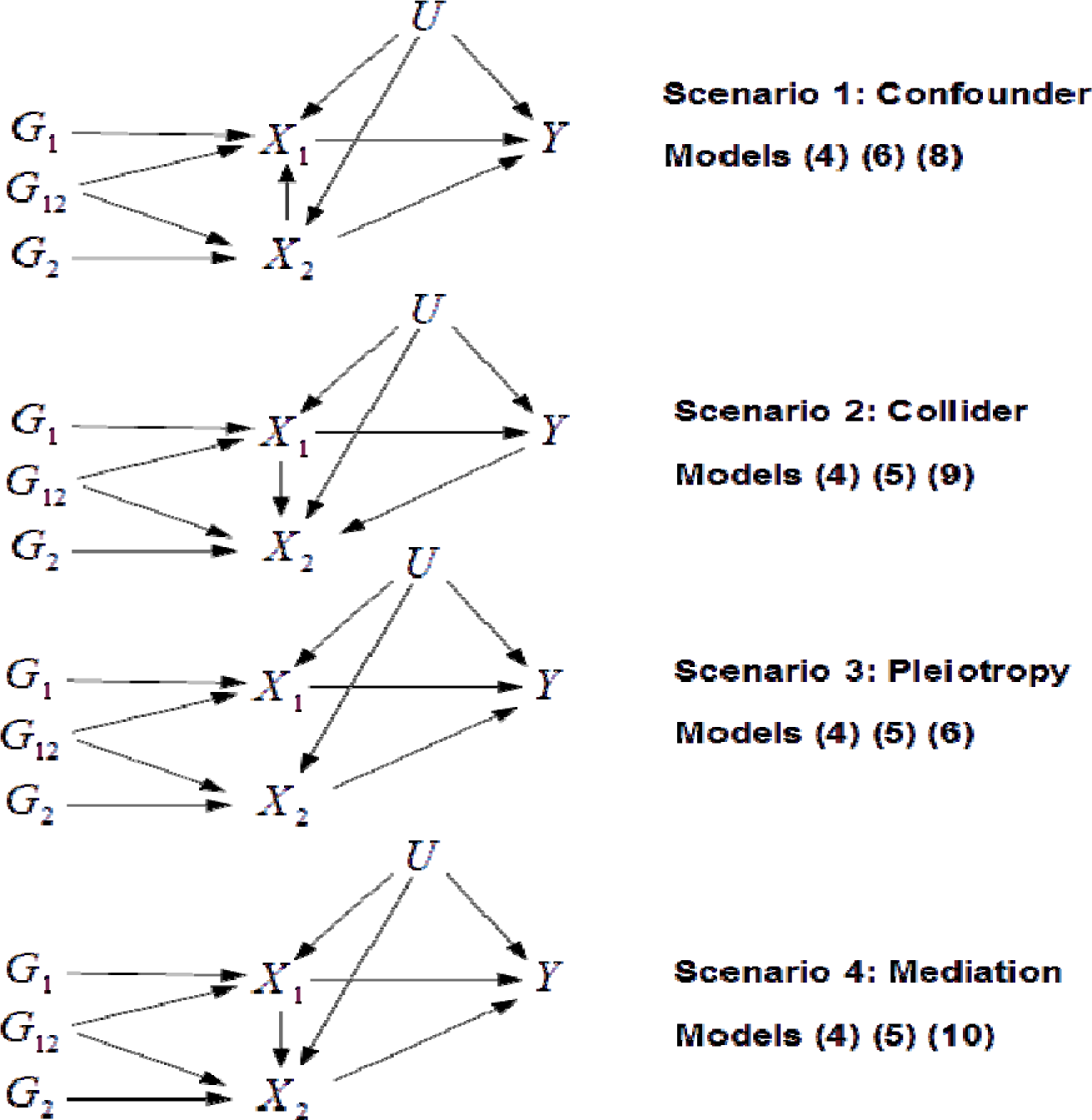
Causal diagrams for scenarios 1-4. Models referred to are the equations above that would give the same relationship between the instruments, exposures and outcome. G_1_, G_2_ and G_12_ are subsets of the full set of SNPs G that affect, and both exposures respectively.

### Simulations

Datasets of 10,000 individuals are simulated under all four scenarios discussed using *L* = 30 genetic variants. The variants are assumed to be uncorrelated but, for added realism and complexity, are further subdivided into three categories:

- 10 SNPs that only predict *X*_1_: *G*_1_ (with a non-zero rr_1_ element but zero rr_2_ element);
- 10 SNPs that only predict *X*_2_: *G*_2_ (with a non-zero rr_2_ element but zero rr_1_ element);
- 10 SNPs that predict *X*_1_ and *X*_2_: *G*_12_ (with non-zero rr_1_ and rr_2_ elements).

*G* therefore represents the complete vector (*G*_1_,*G*_2_,*G*_12_). For each scenario the causal parameter of interest, *β*_1_, is set to 1.

For each scenario, we estimate the causal effects *β*_1_ and *β*_2_ of *X*_1_ and *X*_2_ on *Y*, using a range of estimation methods. With **single sample individual level data**, we implemented:

- OLS, both for *X*_1_ and *X*_2_ individually (i.e. univariable regressions) and together (i.e. a multivariable regression);
- MR for *X*_1_ and *X*_2_ individually, each time using all the available SNPs as instruments,
- Multivariable MR including both *X*_1_ and *X*_2_ in the same analysis.
- MR for *X*_1_ and *X*_2_ individually using only the SNPs that are valid instruments for that exposure (*G*_1_ and *G*_2_ respectively)

With **two sample summary level data**, we implemented:

- MR for *X*_1_ and *X*_2_ individually using all of the instruments available;
- MVMR including both *X*_1_ and *X*_2_;
- MR for *X*_1_ and *X*_2_ individually using only the SNPs that are valid instruments for the exposure.

All estimation methods are described in Table S.1. In all of the scenarios considered the exposure variables are strongly predicted by the instruments and the instruments have no additional pleiotropic effects on the outcome, other than through the exposures included in the model.

### Results

Focusing our attention on exposure *X*_1_, the results from these simulations show that MVMR always gives an unbiased estimate of the direct effect of *X*_1_ on the outcome. In the hypothetical case where only the valid SNPs for *X*_1_ (*G*_1_) are used as instruments in a single variable MR the estimated effect of *X*_1_ on Y is the total effect of a change in *X*_1_ on the outcome. Whether the direct or total effect is of more interest to practitioners will depend on the particular situation being considered. In many of the scenarios explored the direct effect equals the total causal effect, however when *X*_2_ is a mediator of the relationship between *X*_1_ and the outcome, the direct and total effects of *X*_1_ may be substantially different. In this scenario MVMR is not a form of mediation analysis but instead estimates the direct effect of the exposure on the outcome that doesn’t act via the mediator. The results from the simulations are given in Table S.2 and a summary table of what is estimated by each method in each scenario is given in Table 1.

**Table 1.** Summary of estimated effects for *β*_1_

| **Method** | Scenario/which estimand is targeted? |  |  |  |
| --- | --- | --- | --- | --- |
|  | 1 | 2 | 3 | 4 |
| **Individual level data** |  |  |  |  |
| OLS | x | x | x | x |
| Univariate MR | x | Direct/total | x | x |
| MVMR | Direct/total | Direct/total | Direct/total | *Direct* |
| Univariate MR – subset of SNPS | Direct/total | Direct/total | Direct/total | *Total* |
| **Two-sample summary data analysis** |  |  |  |  |
| Univariate MR | x | Direct/total | x | x |
| MVMR | Direct/total | Direct/total | Direct/total | *Direct* |
| Univariate MR – subset of SNPS | Direct/total | Direct/total | Direct/total | *Total* |

When conducting the univariable MR estimation with a subset of the SNPs in *G* we have, for illustration, assumed ‘oracle’ knowledge on which SNPs are valid instruments for each exposure. This will, of course, not be possible in practice For example, in scenario 1 if we select SNPs because they are associated with a *X*_1_ we will select the entire set G, but this will include the subset (*G*_2_,*G*_12_) which exert pleiotropic effects on the outcome Y and thus bias the analysis. Table 1 indeed shows that when all SNPs in G are used for a univariable MR analysis, it will deliver a biased and inconsistent estimate of the total causal effect of *X*_1_ on Y in scenarios 1, 3 and 4. MVMR, by contrast will then provide a consistent estimator of the direct effect of the exposure on the outcome, the consistency of IV analysis under a range of scenarios that include those discussed here has been proved elsewhere. (3, 4, 30) These simulation results also highlight that MVMR does not introduce bias into the results when *X*_2_ is a collider of the relationship between *X*_1_ and *Y*. This is because the predicted value of *X*_2_, *X̂*_2_ which is not dependent on the outcome, is used in the analysis. Of course, adjusting directly for *X*_2_, rather than *X*_2_, would bias the analysis. This is an important benefit of MVMR.

## Testing the assumptions of MVMR

In the simulations above we assumed, for clarity, that the instruments were both strong and valid for the purposes of an MVMR analysis. However, violation of these assumptions can give misleading results in practice, so it is necessary to test these assumptions. We now describe how instrument strength and validity can be scrutinised for an MVMR analysis in the individual and two sample summary data settings.

In addition to assumptions IV1 – IV3 there are additional assumptions and considerations that apply to all instrumental variable estimation, including MR and MVMR. These included the assumptions of linearity and homogeneity which is in many settings required for obtaining a point estimate of the causal effect. Increasing the number of exposures in two sample MVMR will make this a stronger assumption due to the increased number of SNPs and exposures. When implementing MVMR analysis this limitation should be considered and weighed against the benefits when deciding how many exposures to include in the analysis. Another additional assumption, particularly relevant to two-sample MVMR analysis is that all data are drawn from the same underlying population. Throughout our analysis we assume this to hold. The requirement for and issues surrounding this assumption are detailed elsewhere.(31, 32)

### The Individual level data MVMR setting

#### Instrument strength

In any MR analysis the set of genetic instruments G must be strong in order to avoid weak instrument bias (assumption IV1). In single variable MR analysis weak instruments will bias the estimated results in the direction of the observational estimate, however in MVMR analysis it is not clear what direction the bias of the estimation result for each exposure will take as a result of weak instruments.(33) It is therefore important to test the strength of the instruments in any MVMR analysis, however, the assessment of instrument strength is more complicated. It is necessary for G to strongly predict both *X*_1_ and *X*_2_ (as quantified by strong F-statistics), but not sufficient. In addition, G must also jointly predict both *X*_1_ and *X*_2_. That is, once the secondary exposure *X*_2_ has been predicted using *G*, *G* must still be able to predict the primary exposure *X*_1_. Figure 5 illustrates three scenarios (A – C) where this may not be the case even when both exposures appear to be strongly predicted individually by *G* and a fourth scenario (D) where both exposures are strongly predicted.

**Figure 5:**
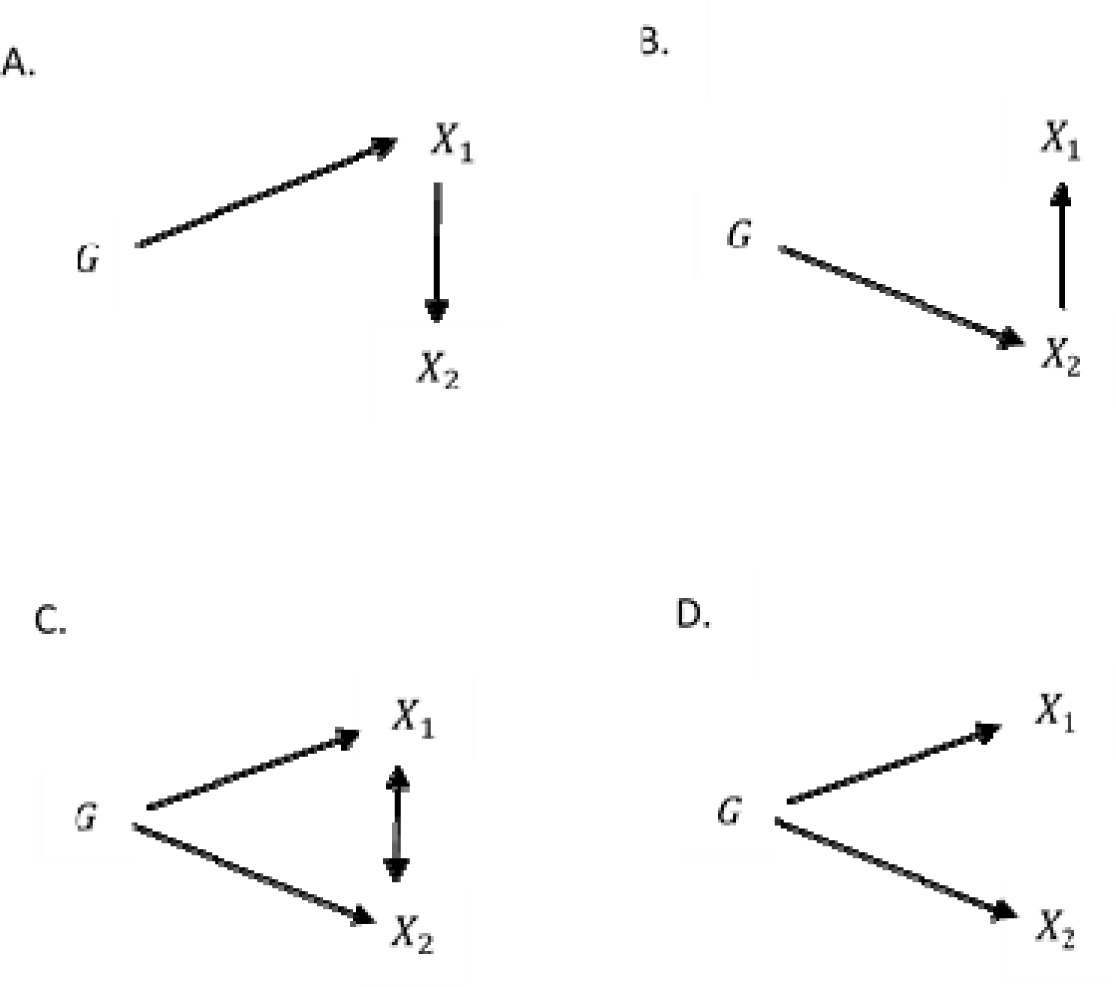
Potential setups of instruments and exposures. In A – B the exposures are individually strongly predicted but are not jointly predicted. In C the exposures are individually strongly predicted but weakly predicted in a joint sense. ․ In D; the exposures are individually and jointly strongly predicted. Specifically: A: G predicts which is a predictor of ․ B: G predicts which is a predictor of ․ C: G predicts and which are highly correlated. D: G predicts and which are uncorrelated (given G).

Joint strength can be assessed using the Sanderson-Windmeijer conditional F statistic(33), , that is available as part of ivreg2 in Stata. is calculated in the following manner:

- is regressed on the full set of genetic instruments (and any control variables included in the estimation) and the predicted value of , is calculated;
- is then regressed on (and any control variables) to yield the TSLS estimate and the residual error terms are saved;
- The errors are then regressed on the full set of instruments (and any control variables). The conditional F statistic is obtained as the F statistic for the effect of the instruments in this regression;
- The conditional F statistic must be adjusted for a degrees of freedom correction, and can be compared to the conventional weak instrument critical values.(34)

For multiple exposure variables the first step is repeated for each of the exposures and all of these predicted values are included in the regression in the second step. This F-statistic can be compared to the standard critical values for weak instruments, therefore if the conditional F statistic for all of the exposure variables are larger than the rule of thumb value of 10 then the instruments can be considered adequately strong for the purposes of MVMR.

#### Instrument validity

If no pleiotropy exists amongst the genetic variants then each one should identify the same causal parameter. This can be evaluated using the Sargan test.(15) Specifically, it tests whether the instruments can explain any of the variation in the outcomes that has not been explained by the value of the exposure variables. It is calculated by the following steps;

- Regress the outcome Y on the exposures using TSLS to yield causal estimates *β̂*_1_ and *β̂*_2_ Calculate the residual error term *Y* - (*β̂*_1_*X*_1_ + *β̂*_2_*X*_2_) and then regress the residuals on the full set of instruments. The Sargan test is then the sample size times the R2 of this regression.
- Evaluating with the Sargan statistic with respect to a *χ*^2^ distribution with degrees of freedom equal to the number of instruments minus the number of predicted exposure variables (i.e. the null hypothesis that all of the instruments are valid).(4)

This test is available as part of the ivreg2 command in Stata, and the ivpack package in R. In order to conduct this test the model must be over-identified, i.e. there must be more instruments than exposure variables (so that the degrees of freedom of the *χ*^2^ test is positive).(35) This ‘global’ test does not give any indication as to which of the genetic instruments are invalid if the test rejects the null. However, alternative methods of estimation can be used to estimate the causal effects as long as at least 50% of the SNPs do not have a pleiotropic effect on the outcome.(36, 37)

### The two-sample summary data setting

Assessment of instrument validity and strength is apparently yet to be described in the two sample summary data setting that is relevant to the majority of contemporary Mendelian randomization studies, and consequently it is not implemented in any standard software. We therefore describe the necessary procedures in fine detail so that they can be confidently implemented by others.

#### Assessing instrument strength: heterogeneity is ‘good’

Suppose that all of the genetic instruments predict both exposure variables, so that models (4), (5) and (6) hold, but there are at least two elements of *π*_1_ and *π*_2_ in (5) and (6) which differ. If true, then the model will be *at least exactly identified*. That is, there will be at least as many independent genetic instruments (i.e. 2) as there are exposure variables to be instrumented. This implies that model (11):

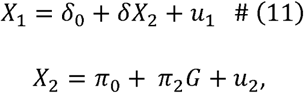

must be over-identified (or equivalently miss-specified), because *X*_2_ cannot then be simply a scalar multiple, *δ*, of *X*_1_ ․ Therefore, we can test for under-identification in our estimation model by testing for over-identification in model (11) using the Sargan test as described above. The equivalence of this test with the Sanderson-Windmeijer approach has been shown formally elsewhere (38) ․ The null of this Sargan test is that of underidentification.

Extending this to two-sample analysis; *π̂*_1,*j*_ =*δπ̂*_2,*j*_ + *ε*_1,*j*_ is analogous to equation (11) estimated by IV using individual level data with *X̂*_2_ predicted using G, therefore it should be possible to test for under-identification in two-sample MVMR estimation by testing for overidentification in the model *π̂*_1,*j*_ = *δπ̂*_2,*j*_ + *v*. We recommend that this test is conducted using a modified version of Cochran’s Q statistic, as shown in equation (12) below:

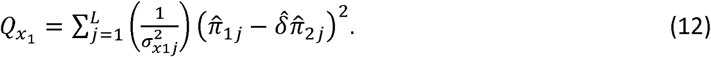

The variance term for 
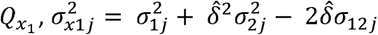
, where 
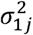
 is the variance of 
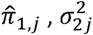
 the variance of *π̂*_2,*j*_, (*σ*_12*j*_ is the covariance of *π̂*_1,*j*_ and *π̂*_2,*j*_, and or is an efficient estimator for *δ* Estimation of the 
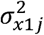
 terms in practice depends on the type of model used to obtain *π̂*_1,*j*_ and *π̂*_2,*j*_. When each exposure is regressed on the entire set of SNPs simultaneously (i.e. via multivariable regressions with an intercept):

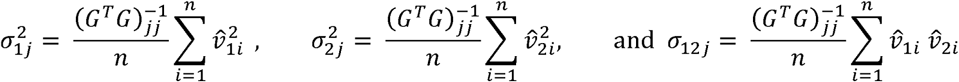

Where *n* is the number of subjects, and (*v̂*_1*i*_,*v̂*_2*i*_) are the estimated residuals from these regressions. If *π̂*_1*j*_ and *π̂*_2*j*_ are obtained separately (i.e. via univariable regressions with an intercept), then the error terms are obtained from the equivalent expressions:

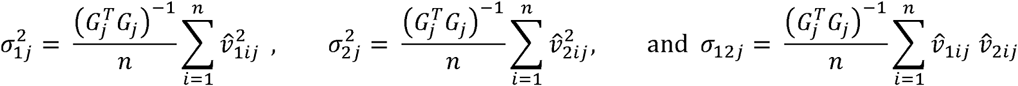

Respectively, *v̂*_1*ij*_ and *v̂*_2*ij*_ are the estimated residuals from the j’th regression.

Under the null hypothesis that the instruments do not contain enough information to predict both exposure variables, *Q*_*x*_1__ will be asymptotically 
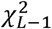
 distributed when *δ* is estimated using an asymptotically efficient estimator, where *L* is the number of instruments. Rejection of the null hypothesis (i.e. detection of ‘heterogeneity’) indicates that the model we wish to estimate *is* identified for *X*_1_.

All the above can be repeated for *X*_2_ by swapping the roles of *π̂*_1_ and *π̂*_2_ and calculating an equivalent *Q* statistic for *X*_2_, *Q_x_2__* say. If both *Q*_*x*_1__ and *Q*_*x*_2__ are larger than the chosen critical value then the null hypothesis of under-identification can be rejected and the test suggests that the instruments can predict variation in both exposures. Table 2 shows the distribution of *Q*_*x*_1__ and *Q*_*x*_2__ for four different scenarios with two exposure variables and *L* = 100 SNPs. *X*_1_ and *X*_2_ are both functions of a set of SNPs and independent confounding variables. In the first simulation the model has been set up as given in Scenario 3 in Figure 4 and in Figure 5D with each of the exposure variables predicted by a set of SNPs and a common confounding variable. This model is identified as both exposure variables can be predicted by the set of instruments. In the second and third simulations the model has been set up in the same way but with no effect of the SNPs on either *X*_1_ or *X*_2_ respectively. That is, the model is underidentified with one of the exposure variables not being predicted by the instruments in each case. In the final simulation the model has been set up with the effect of the SNPs on the exposures as given in Figure 5A and a common confounder. This setup leads to neither exposure being predicted by the SNPs when they are both included in an MVMR estimation as the SNPs in the model cannot predict both of the exposure variables jointly. The results from these simulations show that this test has the required distribution under the null hypothesis.

**Table 2.** The distribution of the modified Q statistic as a test for under-identification

| | $Q_{x_1}$ | | | $Q_{x_2}$ | | |
| --- | --- | --- | --- | --- | --- | --- |
|  | Mean | Std. dev | Rej. Rate (%) | Mean | Std. dev | Rej. Rate (%) |
| $x_1$ strongly identified<br>$x_2$ strongly identified | 1953018 | 26441 | 100 | 1952738 | 247541 | 100 |
| $x_1$ unidentified<br>$x_2$ strongly identified | 99.3 | 14.6 | 6.2 | 379282 | 135506 | 100 |
| $x_1$ strongly identified<br>$x_2$ unidentified | 453058 | 1607136 | 100 | 100.2 | 14.6 | 6.6 |
| $x_1$ strongly identified<br>$x_2$ strongly identified<br>Jointly unidentified<br>$x_1 = \delta x_2, \delta = 1$ | 100.2 | 14.6 | 6.6 | 100.2 | 14.6 | 6.6 |

N = 50,000. Repetitions = 1000, 100 SNPs as instruments. Rejection rates give the proportion of times each $Q$ statistic is larger than the 95<sup>th</sup> percentile of a Chi-squared distribution on 99 degrees of freedom (123.2).

#### Testing instrument validity: heterogeneity is ‘bad’

Cochran’s Q statistic for the regression of interest has been proposed as a method for identifying the presence of invalid instruments (e.g. due to horizontal pleiotropy) in two-sample summary data MR analysis, with a single exposure.(19) Specifically, if all instruments are valid IVs, and the modelling assumptions necessary for two-sample MR are satisfied, then each genetic instrument should give the same estimate of the effect of the exposure on the outcome. Excessive heterogeneity in the causal effect estimates obtained by each SNP individually now becomes an indicator of invalid instruments. We propose testing for invalidity in two sample summary data MVMR using an adjusted version of the Cochran Q statistic given by:

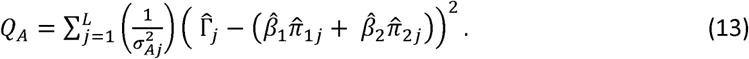

Where 
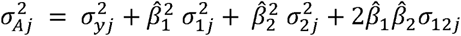
. To clarify, 
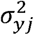
 is the variance of Γ̂_j_, and *β̂*_1_ and *β̂*_2_ are efficient estimates of *β*_1_ and *β*_2_ (for example as obtained from fitting model (7)). Under the null hypothesis that the genetic instruments do not have pleiotropic effects on the outcome, *Q_A_* is asymptotically *χ^2^* distributed with (*L* − 2) degrees of freedom. The standard implementation of Cochran’s Q would merely have a weighting of 
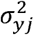
, and is not therefore asymptotically *χ^2^* distributed. It is a straightforward generalisation of the adjusted Q statistic recently proposed by Bowden et al in the univariable MR setting.(18) Excessive heterogeneity in *Q_A_* therefore brings assumptions IV2 and IV3 into doubt.

Figure 6 shows the distribution of *Q_A_* compared to the standard Q statistic and a *X*^2^ distribution with 98 degrees of freedom for a model with 2 exposure variables and 100 genetic instruments. For simplicity the estimated effects of the SNPs on the exposures each have a common variance of 0.02 and have a common covariance of 0. *Q_A_* is seen to have the correct distribution under the null hypothesis of no pleiotropy in the model.

**Figure 6:**
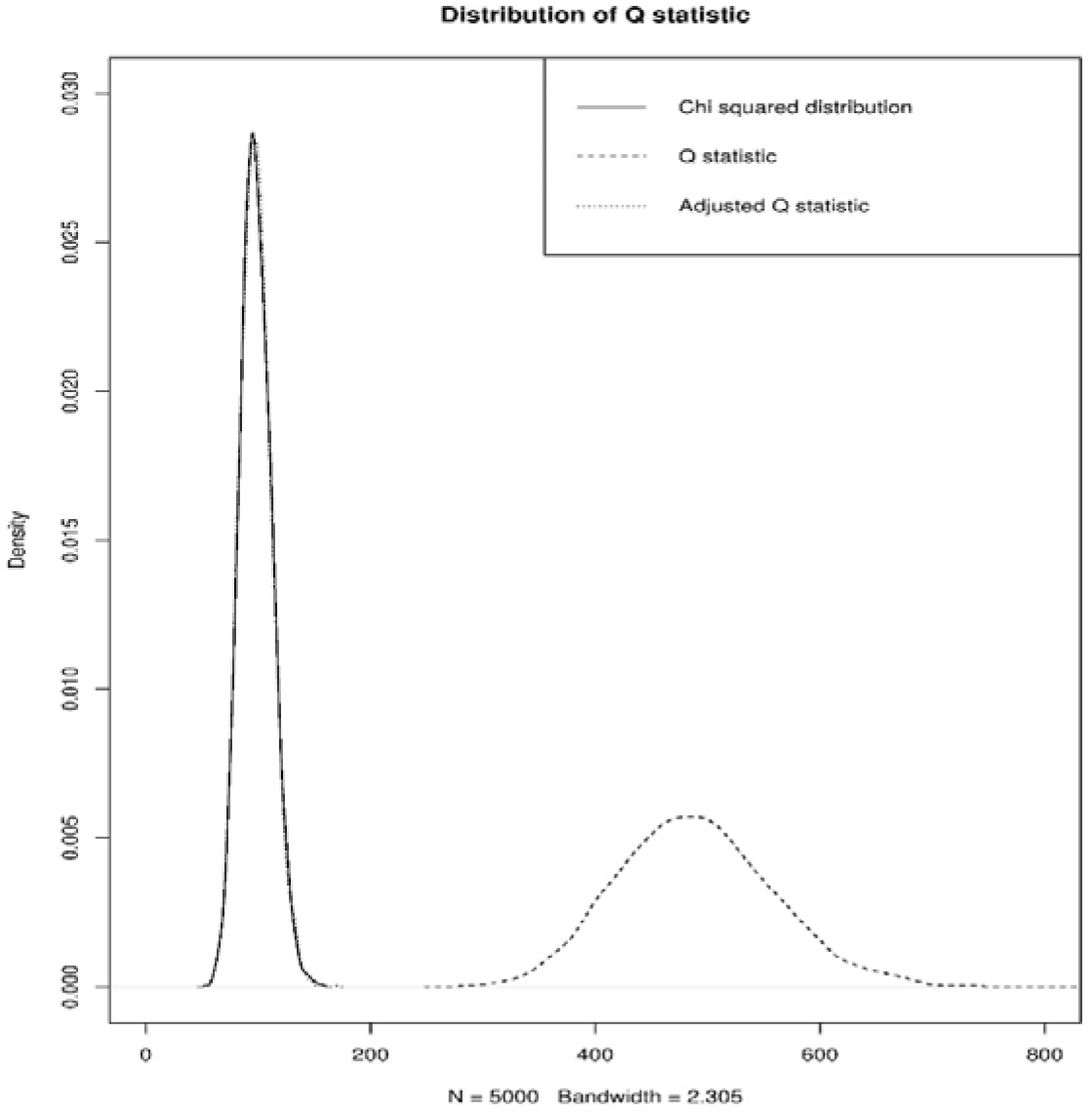
The distribution of the adjusted and standard Q statistics under the null hypothesis of no heterogeneity. 5000 repetitions, 100 SNPs. Here *β*_1_ = *β*_2_ = 1. 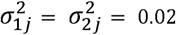, *σ*_12*j*_ = 0 for all j.

We suggest updating the two sample causal estimates in an iterative process using weights derived from the initial estimates of the causal effects, which is referred to as ‘modified iterative’ weighting in Bowden et al (18) within the context of a univariate MR analysis. Further work is required to fully investigate the effect of this and to understand how the fully analytical solution discussed in (18) which finds the causal estimate that directly minimises an equivalent Q statistic, could be extended to the multivariable case, but if done so this could help to mitigate the effect of weak instrument bias.

### Approximating *Q*_*x*1_, *Q*_*x*2_ and *Q_A_* with incomplete information

The covariance vector *σ*_12*j*_ that is necessary for correct specification of *Q*_*x*1_, *Q*_*x*2_ and *Q_A_* can only be calculated from the individual participant data. If this information is not available, one solution would be to ensure that *σ_12j_* is zero, by estimating the genetic associations with each exposure and the outcome in separate samples. This would correspond to a ‘three-sample’ summary data MR-analysis when two exposures constitute the MVMR analysis.

Another pragmatic solution would be to assume that each *π_12j_* term is zero. This will give a good approximation for *Q*_*x*1_ and *Q*_*x*2_ whenever *δσ_12j_* is small and for *Q_A_* whenever *β̂*_1_*β̂*_2_*σ*_12j_ is small.

## Application to education, cognitive ability and Body Mass Index

In this section we apply the methods discussed above to investigate whether there is evidence for a causal effect of education and cognitive ability on body mass index (BMI) using data from UK biobank. Education and cognitive ability have both been found to be associated with BMI, with higher levels of education and cognitive ability being associated with lower levels of BMI.(39–42) However, there is also a high level of correlation observed between completed education and measured cognitive ability, therefore it is not clear whether, once this correlation has been controlled for, both education and cognitive ability have a causal effect on BMI.(39)

### Data

UK biobank recruited 502,641 individuals aged 37-73 years between 2006 and 2010 from across the UK. Individuals where invited to a clinic where they answered a questionnaire and interview about a range of health topics and provided anthropomorphic measurements and gave samples of blood, urine and saliva. This study has been described in full previously.(43)

Individuals in UK biobank were asked to report the highest educational qualification they had obtained. For each individual we assigned an age at which they left education based their reported qualification. A breakdown of educational qualifications and associated ages across the cohort is given in Table S.3.

Cognitive ability was measured among a subset of the UK biobank participants as the number of correct answers recorded in a series of 13 questions designed to measure cognitive ability that where completed as part of the initial clinic. The cognitive ability variable was then standardised to have mean zero and variance 1. BMI was calculated based on the height and weight of the individuals in the sample. Throughout the analysis we analysed this variable on the natural log scale because of its skewed distribution.

### Analysis

We first conducted MR analyses for the effect of education and cognitive ability on BMI separately using single variable MR. A single composite instrument for education was created using the polygenic score of 74 SNPs from a recent GWAS of educational attainment.(44) A single composite instrument for cognitive ability was created using the polygenic score of 18 SNPs from a recent GWAS of cognition.(45) As this GWAS was conducted using the interim release of UK Biobank we restricted our analysis to individuals not included in the interim release.

We then conducted a multivariable MR analysis of the effect of education and cognitive ability on BMI. This analysis included both the composite instruments for education and cognitive ability used in the single variable MR analyses.

The results from this analyses, along with a multivariable OLS regression of BMI on education and cognitive ability, are given in Table 3. The OLS results show that each extra year of education is associated with a decrease in BMI MR and MVMR results suggests a causal effect in the same direction, but with a larger magnitude. The results for cognitive ability are more mixed with no association seen in the OLS results, a negative total effect of cognitive ability on BMI in the MR analysis and potentially a positive direct effect of cognitive ability on BMI observed in the MVMR analysis. Our empirical and theoretical investigation helps to clarify why the the high level of correlation between education and cognitive ability would lead to the conclusion that there is a negative effect of cognitive ability on BMI in MR analysis. The MVMR results show that, if anything, the direct effect of increasing cogntive ability is to increase BMI. These results highlight the potential benefits of MVMR. However, before giving much credence to this result it is necessary to assess the strength of our SNPs to jointly predict education and cognitive ability.

**Table 3.** The effect of education and cognitive ability on BMI

|  |  | OLS |  | MR |  |
| --- | --- | --- | --- | --- | --- |
|  |  | Single variable | Multivariable | Single variable | Multivariable |
| **Age completed Education** | *Effect* | -0.008 | -0.008 | -0.028 | -0.044 |
|  | *Std.Error* | (0.0003) | (0.0003) | 0.005 | 0.013 |
|  | *95% C.I.* | [-0.0095, -0.0074] | [-0.0085, -0.0074] | [-.0391 - .0179] | [-.0704 - .0187] |
|  | *F-statistic* |  |  | 188.2 | 195.0 |
|  | *S-W F-statistic* |  |  |  | 35.7 |
| **Standardised cognitive ability Score** | *Effect* | -0.006 | 0.0001 |  | 0.048 |
|  | *Std.Error* | (0.0007) | (0.0007) | 0.008 | 0.025 |
|  | *95% C.I.* | [-0.0078, -0.0051] | [-0.0013, 0.0014] | [-.0380 - .0082] | [-0.001 0.098] |
|  | *F-statistic* |  |  | 542.2 | 309.7 |
|  | *S-W F-statistic* |  |  |  | 37.0 |

Dependent variable is log(BMI).
Estimates of the effect of education and cognitive ability on BMI from OLS, single variable MR and multivariable MR analysis of individual level data.
All regressions also include a full set of control variables: age, gender, income and 10 genetic principal components. Instruments are constructed from GWAS scores for education and cognitive ability. The regressions are weighted so that individuals who left school at 15 are given an 80% upweighting. All non-European and related individuals have been excluded from the analysis. Total sample size included in all regressions: 74,309.

### Testing the instrument strength in the single sample setting

As a measure of the strength of the instruments we calculate the standard F-statistic for both education and cognitive ability and the Sanderson – Windmeijer partial F-statstic(33) for the multivariable MR analysis. As all F-statistics are much larger than the rule-of-thumb cut off of 10 we are reassured that the instruments are not individually weak. However, the partial F-statistic for both education and cognitive ability is significantly lower, showing that the power of the instruments to predict both variables simultanously is greatly reduced.

The Sargan test for invalid instruments can only be calculated for estimation models with more instruments than exposure variables. In this estimation we have two exposure and two instruments and so it is not possible to calculate the Sargan statistic.

### Two-Sample Multivariable MR

To illustrate two-sample MVMR we randomly divided the sample used for the individual analysis into three equal-sized groups. For each SNP used in the polygenic score, we then calculated its effect on log(BMI), education and cognitive ability using different parts of the sample., The results were then used to conduct a two-sample MVMR analysis. The results are given in Table S.4. They show that increased education has a direct effect which decreases BMI and cognitive ability has no direct effect on BMI. The results are in line with those obtained from the individual level analysis.

### Testing instrument strength in the two-sample setting

To test for weak instruments in this analysis we have calculated the weak-instrument *Q* statistics for education and cognitive ability. The *Q_edu_* statistic for education is 1724.4. The *Q_cog_* statistic for cognitive ability is 1488.8. The critical value for a *χ*^2^ distribution with 88 degrees of freedom at the 5% level is 110.9. Therefore we reject the null hypothesis that the SNPs do not explain any of the variation in the exposures education and cognitive ability in this two sample analysis and can conclude that these SNPs can predict both education and cognitive ability in the data.

### Testing for pleiotropy in the two-sample setting

To illustrate the two tests for pleiotropy discussed earlier we report the *Q_A_* statistic for MVMR. The value of *Q_A_* for this regression is 129.5. The critical value for a *χ*^2^ distribution with 87 degrees of freedom is 109.77. Therefore, the null hypothesis that there is no heterogeneity is rejected for this value of *Q_A_*.

### Multivariable MR Egger regression

An alternative procedure that has been recently proposed to adjust for pleiotropy beyond that explainable by genetically predictable exposures (e.g. *X*_1_ and *X*_2_) is a Multivariable MR Egger regression(46) This is a natural extension of the original MR Egger approach(1) and is calculated by fitting the two sample MVMR model with a constant included;

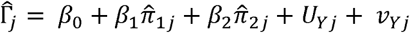

If the constant is different from zero this suggests that additional pleiotropy is meaningfully biasing the analysis. However a generalisation of the InSIDE assumption is required in order for it to deliver unbiased causal estimates. These are described in detail elsewhere.(1)

The two sample results were used to fit multivariable MR Egger regression, the results of which are given in Table S.5. Its intercept parameter is estimated to be small, and consequently the estimated effects of the exposures do not differ from those in the two-sample MVMR estimation. This supports the suggestion that the SNPs do not exert a direct effect on BMI apart from through education or cognitive ability. As MR-Egger is dependent on the orientation of the SNP exposure associations, we repeated this analysis with the associations orientated so that the SNP education associations where all positive and then with the SNP cognitive ability associations all positive. These changes had no substantive effect on the results obtained.

The difference between the Q-statistic and Multivariable MR Egger estimation suggest an inconsistency between these two tests however this may have arisen due to a high level of variation in the effect of the SNPs on each exposure leading to a higher Q statistic. This is supported by Figure. 7a and 7b which gives individual MR plots for each exposure, and shows that there is a large amount of variation of the SNPs on each of the exposures. Repeating this analysis with the outlying SNP excluded makes no substantive difference to the results obtained.

**Figure 7:**
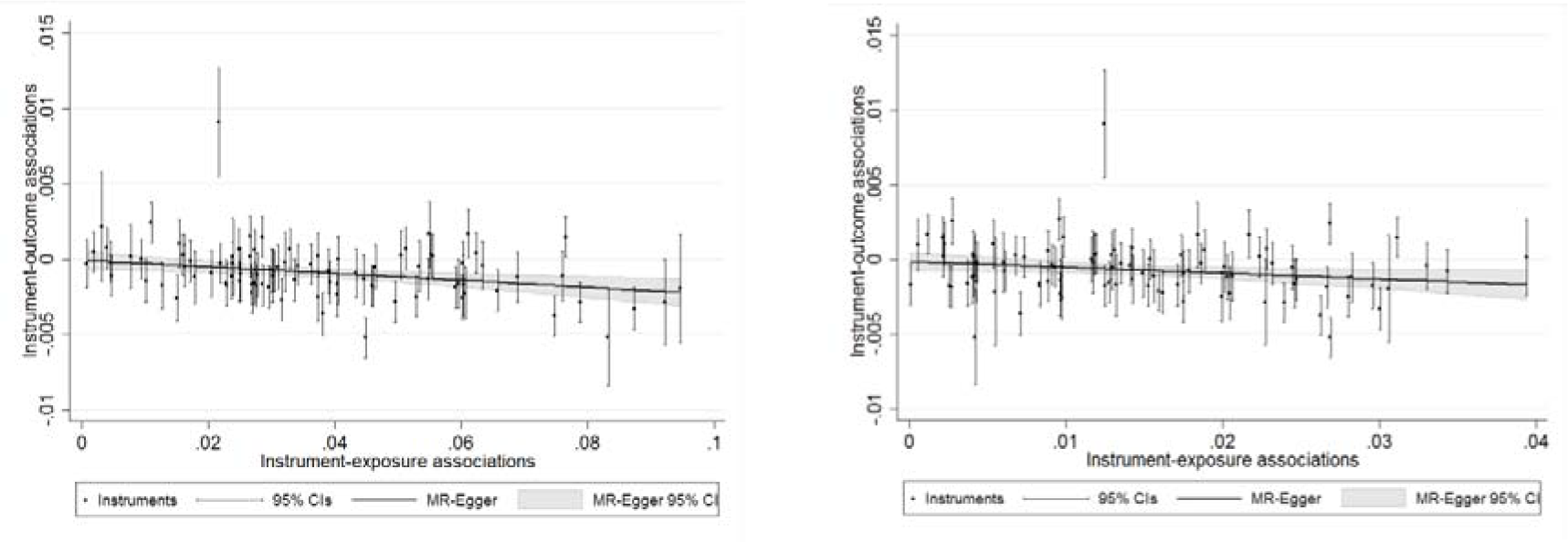
Left: MR Egger plot for the association between educational attainment and BMI. Right: MR Egger plot for the association between cognitive ability and BMI. All SNPs that affect either education or cognitive ability are included

The MVMR Egger analysis was repeated using the effect of each SNP on education, cognitive ability and BMI taken from GWAS estimates.(44, 45, 47) The magnitude of the estimated effects differ in this analysis as the outcome variable is BMI rather than the natural log of BMI, however these results also show no pleiotropic effect of the SNPs on the outcome and a negative effect of higher education on BMI. Results from this analysis are given in Table S.5.

## Discussion

In this paper we have attempted to explain the principled application and interpretation of instrumental variable analysis to the epidemiological setting with multiple exposures. We first focused on the individual data setting, for which it is possible to borrow well-established methods (and related software) from the econometrics literature. We then considered the two-sample setting and built upon previous research in this area by developing new tests for assessing the validity and relevance of the genetic instruments. In particular, we propose two new tests;

- Modified Q statistics, (in our case, and) for instrument relevance that detect ‘good’ heterogeneity if a set of SNPs can jointly and reliably predict all intermediate exposures of interest;
- A modified Q statistic, for instrument validity that detects ‘bad’ heterogeneity if a set of SNPs contains invalid instruments.

We finally illustrated the application of MVMR using individual and summary level data to estimate the effect of education and cognitive ability on BMI. The results from this analysis show that increasing education leads to lower BMI and the size of this effect increases when cognitive ability is controlled for. Comparing the single exposure MR analysis results (with all SNPs that affect educational attainment excluded) to the MVMR results for cognitive ability shows a large change in the size and direction of the effect. This result suggests that education is a mediator of the relationship between cognitive ability and BMI and any direct effect of cognitive ability on BMI is minimal.

The methods we describe can be used to estimate the effect of multiple related exposures on an outcome using either individual level or summary level data. Although we have focused on the case of two exposures for ease of explanation, all of these methods can be easily applied to scenarios with three or more exposures. An advantage of MVMR analysis is that SNPs which are thought to potentially affect multiple exposure variables, or where it is not clear exactly which exposure they affect, can be included when estimating the effects of the exposures on the outcome. This makes MVMR particularly useful when the exposures are closely related or one (or more) is thought to be a potential pleiotropic pathway from the SNPs to the outcome. MVMR will also produce consistent estimates when there is measurement error in any of the exposure variables and therefore is a useful method of analysis when multiple exposure variables are thought to be subject to measurement error.

As with all MR analysis it is important to ensure that the IV assumptions are satisfied. Here we explain how the IV assumptions apply to a MVMR analysis. We describe existing tests that can be used to test the assumptions in individual level data and propose tests that can be used with two-sample summary level data. These new tests are a key strength of this work as MVMR cannot be effectively used as part of the tools a researcher has available for analysis unless the potential pitfalls of the analysis are well understood. Our applied results highlight the importance of considering the IV assumptions in the context of the particular analysis being conducted as even when the instruments appear to be very strong for each of the exposures individually, this does not guarantee that they will be equally as strong for the exposures when estimated jointly in a MVMR model. For example, the F-statistics decrease from 195 and 310 to 36 and 37 for educational attainment and cognitive ability respectively.

A practical limitation of the new tests we develop for two-sample summary data MVMR is the reliance on knowledge of the covariance between the effect of the SNP on each exposure. These results are not available in conventional GWAS results, and it would be infeasible to calculate them in advance for every possible combination of exposure variable that could be included in a MVMR model. Unfortunately, our work shows that this information is strictly needed for valid inference. In order to conduct these tests in summary level data we therefore have to make a choice about how to treat these missing pieces of information. If the data were available it can be directly calculated from the individual level data for the particular MVMR study being conducted. Alternatively, it could be assumed to be zero, or set to zero by using non-overlapping GWAS studies for each exposure as the standard error of the estimated SNP effects will not correlate across different samples. This is an important limitation of the results given here for testing the assumptions of two-sample summary data MVMR.

Another weakness of the instrument relevance test we develop is that this is a test for whether the SNPs can conditionally explain any of the variation in the exposure variables, rather than being a more usual weak instrument test, such as the rule of thumb of F being greater than 10 for a univariable MR analysis or the Sanderson-Windmeijer conditional F statistic for IV analysis with individual level data. Extending this test to weak instrument is an area for future work.

## Acknowledgements

This research was supported by the UK Medical Research Council Integrative Epidemiology Unit at the University of Bristol (MC_UU_00011/2, MC_UU_00011/1). The analysis has been conducted using the UK Biobank Resource as part of application 8786.

## References

1. Bowden J, Smith GD, Burgess S. Mendelian randomization with invalid instruments: effect estimation and bias detection through Egger regression. Int J Epidemiol. 2015;44:512–25.

2. Burgess S, Dudbridge F, Thompson SG. Re:”Multivariable Mendelian randomization: the use of pleiotropic genetic variants to estimate causal effects”. Am J Epidemiol. 2015;181(4):290–1.

3. Davidson R, MacKinnon JG. Estimation and inference in econometrics. OUP Catalogue. 1993.

4. Wooldridge JM. Econometric analysis of cross section and panel data: MIT press; 2010.

5. Thompson John R, Minelli C, Del Greco MF. Mendelian Randomization using Public Data from Genetic Consortia. Int J Biostat 2016.

6. Wald A. The fitting of straight lines if both variables are subject to error. Ann Stat. 1940;11(3):284–300.

7. Burgess S, Bowden J. Integrating summarized data from multiple genetic variants in Mendelian randomization: bias and coverage properties of inverse-variance weighted methods. arXiv preprint arXiv:151204486. 2015.

8. Lawlor DA, Harbord RM, Sterne JA, Timpson N, Davey Smith G. Mendelian randomization: using genes as instruments for making causal inferences in epidemiology. Stat Med. 2008;27(8):1133–63.

9. Davey Smith G, Ebrahim S. ‘Mendelian randomization’: can genetic epidemiology contribute to understanding environmental determinants of disease?*. Int J Epidemiol. 2003;32(1):1–22.

10. Davey Smith G, Hemani G. Mendelian randomization: genetic anchors for causal inference in epidemiological studies. Hum Mol Genet. 2014;23(R1):R89–R98.

11. Burgess S, Thompson SG. Avoiding bias from weak instruments in Mendelian randomization studies. Int J Epidemiol. 2011;40(3):755–64.

12. Bennett DA, Holmes MV. Mendelian randomisation in cardiovascular research: an introduction for clinicians. Heart. 2017:heartjnl-2016-310605.

13. Haycock PC, Burgess S, Wade KH, Bowden J, Relton C, Smith GD. Best (but oft-forgotten) practices: the design, analysis, and interpretation of Mendelian randomization studies. Am J Clin Nutr. 2016;103(4):965–78.

14. VanderWeele TJ, Tchetgen EJT, Cornelis M, Kraft P. Methodological challenges in Mendelian randomization. Epidemiology (Cambridge, Mass). 2014;25(3):427.

15. Sargan JD. The Estimation of Economic Relationships using Instrumental Variables. Econometrica. 1958;26(3):393–415.

16. Windmeijer F. Two-Stage Least Squares as Minimum Distance. Econom J. 2018.

17. Cochran WG. The combination of estimates from different experiments. Biometrics. 1954;10(1):101–29.

18. Bowden J, Fabiola Del Greco M, Minelli C, Lawlor D, Sheehan N, Thompson J, et al. Improving the accuracy of two-sample summary data Mendelian randomization: moving beyond the NOME assumption. bioRxiv. 2017:159442.

19. Greco M, Del F, Minelli C, Sheehan NA, Thompson JR. Detecting pleiotropy in Mendelian randomisation studies with summary data and a continuous outcome. Stat Med. 2015;34(21):2926–40.

20. Bowden J, Del Greco MF, Minelli C, Davey Smith G, Sheehan N, Thompson J. A framework for the investigation of pleiotropy in two-sample summary data Mendelian randomization. Stat Med. 2017;36(11):1783–802.

21. Burgess S, Dudbridge F, Thompson SG. Combining information on multiple instrumental variables in Mendelian randomization: comparison of allele score and summarized data methods. Stat Med. 2016;35(11):1880–906.

22. Burgess S, Butterworth AS, Thompson JR. Beyond Mendelian randomization: how to interpret evidence of shared genetic predictors. J Clin Epidemiol. 2016;69:208–16.

23. Swanson SA, Labrecque J, Hernán MA. Causal null hypotheses of sustained treatment strategies: What can be tested with an instrumental variable? Eur J Epidemiol. 2018:1–6.

24. Didelez V, Sheehan N. Mendelian randomization as an instrumental variable approach to causal inference. Stat Methods Med Res. 2007;16(4):309–30.

25. Burgess S, Thompson SG. Use of allele scores as instrumental variables for Mendelian randomization. Int J Epidemiol. 2013;42(4):1134–44.

26. Burgess S, Thompson DJ, Rees JMB, Day FR, Perry JR, Ong KK. Dissecting Causal Pathways Using Mendelian Randomization with Summarized Genetic Data: Application to Age at Menarche and Risk of Breast Cancer. Genetics. 2017.

27. Relton CL, Davey Smith G. Two-step epigenetic Mendelian randomization: a strategy for establishing the causal role of epigenetic processes in pathways to disease. Int J Epidemiol. 2012;41(1):161–76.

28. Munafò MR, Tilling K, Taylor AE, Evans DM, Davey Smith G. Collider scope: when selection bias can substantially influence observed associations. Int J Epidemiol. 2017.

29. Hernán MA, Hernández-Díaz S, Robins JM. A structural approach to selection bias. Epidemiology. 2004:615–25.

30. Bareinboim E, Tian J, Pearl J, editors. Recovering from Selection Bias in Causal and Statistical Inference. AAAI; 2014.

31. Bareinboim E, Pearl J. Causal inference and the data-fusion problem. Proceedings of the National Academy of Sciences. 2016;113(27):7345–52.

32. Hernán MA, Robins JM. Instruments for causal inference: an epidemiologist’s dream? Epidemiology. 2006:360–72.

33. Sanderson E, Windmeijer F. A weak instrument F-test in linear IV models with multiple endogenous variables. J Econom. 2016;190(2):212–21.

34. Stock J, Yogo M. Testing for Weak Instruments in Linear IV Regression. In: Andrews DWK, editor. Identification and Inference for Econometric Models. New York: Cambridge University Press; 2005. p. 80–108.

35. Murray MP. Avoiding invalid instruments and coping with weak instruments. J Econ Perspect. 2006;20(4):111–32.

36. Kang H, Zhang A, Cai TT, Small DS. Instrumental Variables Estimation With Some Invalid Instruments and its Application to Mendelian Randomization. Journal of the American Statistical Association. 2016;111(513):132–44.

37. Windmeijer F, Farbmacher H, Davies N, Smith GD. On the Use of the Lasso for Instrumental Variables Estimation with Some Invalid Instruments. Journal of the American Statistical Association. 2018:1–32.

38. Windmeijer F. Testing Over and Underidentification in Linear Models, with Applications to Dynamic Panel Data and Asset-Pricing Models. University of Bristol Department of Economics Working Paper. 2018.

39. Chandola T, Deary IJ, Blane D, Batty GD. Childhood IQ in relation to obesity and weight gain in adult life: the National Child Development (1958) Study. Int J Obes. 2006;30(9):1422.

40. Benson R, von Hippel PT, Lynch JL. Does more education cause lower BMI, or do lower-BMI individuals become more educated? Evidence from the National Longitudinal Survey of Youth 1979. Soc Sci Med. 2017.

41. Johnson W, Kyvik KO, Skytthe A, Deary IJ, Sørensen TIA. Education Modifies Genetic and Environmental Influences on BMI. PLoS One. 2011;6(1):e16290.

42. Spasojević J. Chapter 9 Effects of Education on Adult Health in Sweden: Results from a Natural Experiment. Current Issues in Health Economics. p. 179–99.

43. Collins R. What makes UK Biobank special? The Lancet. 2012;379(9822):1173–4.

44. Okbay A, Beauchamp JP, Fontana MA, Lee JJ, Pers TH, Rietveld CA, et al. Genome-wide association study identifies 74 loci associated with educational attainment. Nature. 2016;533(7604):539.

45. Sniekers S, Stringer S, Watanabe K, Jansen PR, Coleman JR, Krapohl E, et al. Genome-wide association meta-analysis of 78,308 individuals identifies new loci and genes influencing human intelligence. Nat Genet. 2017.

46. Rees J, Wood A, Burgess S. Extending the MR-Egger method for multivariable Mendelian randomization to correct for both measured and unmeasured pleiotropy. Stat Med. 2017.

47. Speliotes EK, Willer CJ, Berndt SI, Monda KL, Thorleifsson G, Jackson AU, et al. Association analyses of 249,796 individuals reveal 18 new loci associated with body mass index. Nat Genet. 2010;42(11):937–48.

